# QTQTN motif upstream of the furin-cleavage site plays key role in SARS-CoV-2 infection and pathogenesis

**DOI:** 10.1101/2021.12.15.472450

**Authors:** Michelle N. Vu, Kumari G. Lokugamage, Jessica A. Plante, Dionna Scharton, Bryan A. Johnson, Stephanea Sotcheff, Daniele M. Swetnam, Craig Schindewolf, R. Elias Alvarado, Patricia A. Crocquet-Valdes, Kari Debbink, Scott C. Weaver, David H. Walker, Andrew L. Routh, Kenneth S. Plante, Vineet D. Menachery

## Abstract

The furin cleavage site (FCS), an unusual feature in the SARS-CoV-2 spike protein, has been spotlighted as a factor key to facilitating infection and pathogenesis by increasing spike processing ^1,2^. Similarly, the QTQTN motif directly upstream of the FCS is also an unusual feature for group 2B coronaviruses (CoVs). The QTQTN deletion has consistently been observed in *in vitro* cultured virus stocks and some clinical isolates ^3^. To determine whether the QTQTN motif is critical to SARS-CoV-2 replication and pathogenesis, we generated a mutant deleting the QTQTN motif (ΔQTQTN). Here we report that the QTQTN deletion attenuates viral replication in respiratory cells in vitro and attenuates disease *in vivo*. The deletion results in a shortened, more rigid peptide loop that contains the FCS, and is less accessible to host proteases, such as TMPRSS2. Thus, the deletion reduced the efficiency of spike processing and attenuates SARS-CoV-2 infection. Importantly, the QTQTN motif also contains residues that are glycosylated^4^, and disruption its glycosylation also attenuates virus replication in a TMPRSS2-dependent manner. Together, our results reveal that three aspects of the S1/S2 cleavage site – the FCS, loop length, and glycosylation – are required for efficient SARS-CoV-2 replication and pathogenesis.

## Introduction

SARS-CoV-2 emerged in late 2019 and has caused the largest pandemic since the 1918 influenza outbreak ^5^. An unusual feature of SARS-CoV-2 is the presence of a furin cleavage site in its spike protein ^6^. The CoV spike, a trimer of spike proteins composed of the S1 and S2 subunits, responsible for receptor binding and membrane fusion, respectively ^5^. After receptor binding, the spike protein is proteolytically cleaved at the S1 and S2 sites to activate the fusion machinery. For SARS-CoV-2, the spike protein contains a novel cleavage motif recognized by the host cell furin protease (PRRAR) directly upstream of the S1 cleavage site that facilitates cleavage prior to virion release from the producer cell. This furin cleavage site (FCS), not found in other group 2B CoVs, plays a key role in spike processing, infectivity, and pathogenesis as shown by our group and others ^2,7^.

Importantly, another novel amino acid motif, QTQTN, is found directly upstream of the FCS. This QTQTN motif, also absent in other group 2B CoVs, is often deleted and has been pervasive in cultured virus stocks of the alpha, beta, and delta variants ^8^. In addition, the QTQTN deletion has been observed in a small subset of patient samples as well ^9^. Because this deletion has been frequently identified, we set out to characterize it and determine whether it has consequences for viral replication and virulence. Using our infectious clone ^10,11^, we demonstrated that the loss of this motif attenuates SARS-CoV-2 replication in respiratory cells *in vitro* and pathogenesis in hamsters. The QTQTN deletion results in reduced spike cleavage and diminished capacity to use serine proteases on the cell surface for entry. Importantly, mutations of glycosylation-enabling residues in the QTQTN motif results in similar replication attenuation despite intact spike processing. Together, our results highlight elements in the SARS-CoV-2 spike in addition to the furin cleavage site that contribute to increased replication and pathogenesis.

## Results

### ΔQTQTN attenuates viral replication

In addition to the furin cleavage site (FCS), comparison of group 2B coronavirus sequences also revealed the presence of an upstream QTQTN motif directly in the SARS-CoV-2 spike protein; this motif is absent in other coronaviruses except for the closely related RaTG13 bat coronavirus (**Fig. 1a**). Importantly, this QTQTN motif is often deleted in SARS-CoV-2 strains propagated in Vero E6 cells ^8^. To explore the role of the QTQTN motif in SARS-CoV-2 infection and pathogenesis, we generated a mutant in the WA-1 background (early U.S. case from 2020) by deleting QTQTN (ΔQTQTN) using our reverse genetics system ^10,11^ (**Fig. 1b**). Examining the deletion on the SARS-CoV-2 spike structure, our modeling suggested that the ΔQTQTN mutant forms a stable α-helix in the loop containing the S1’ cleavage site (**Fig. 1c**). While the mutant retains the furin cleavage motif, its a-helix is predicted to make the loop less flexible and reduce access to the proteolytic cleavage site.

**Figure 1:**
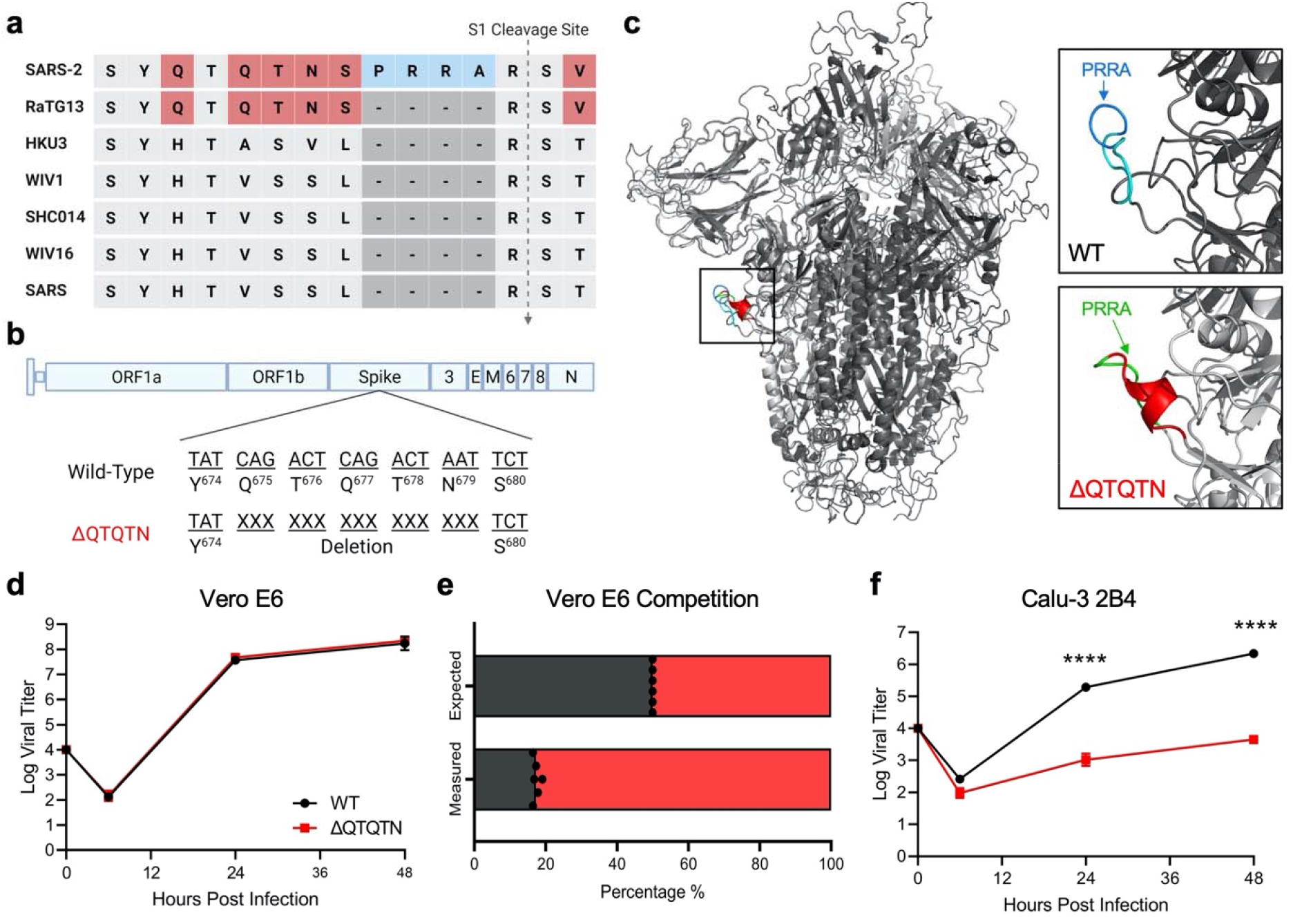
*In vitro* characterization of SARS-CoV-2 ΔQTQTN. **a**, Comparison of S1/S2 cleavage site across SARS-CoV, SARS-CoV-2 and 5 related bat CoVs. **b**, Schematic of SARS-CoV-2 genome with deletion of QTQTN codons. **c**, SARS-CoV-2 spike trimer (grey) with WT (upper) and model-predicted ΔQTQTN (lower) overlaid. PRRA (blue) is exposed with QTQTN (cyan) present in WT and extending the loop (upper). An α-helix is formed with deletion of QTQTN (red) and PRRA (green) is exposed (lower). **d**, Viral titer from Vero E6 cells infected with WT (black) or ΔQTQTN (red) SARS-CoV-2 at an MOI of 0.01 (n=3). **e**, Competition assay between WT and ΔQTQTN SARS-CoV-2 at a ratio of 1:1, showing RNA percentage from next generation sequencing. **f**, Viral titer from Calu-3 2B4 infected with WT or ΔQTQTN SARS-CoV-2 at an MOI of 0.01 (n=3). Data are mean ± s.d. Statistical analysis measured by two-tailed Student’s *t*-test. *, p≤0.05; **, p≤0.01; ***, p≤0.001; ****, p≤0.0001.

The deletion of QTQTN motif did not affect virus replication in Vero E6 (African green monkey kidney cells) cells with the rescue stock titer comparable to wild-type WA-1 (WT) in yield; yet, the ΔQTQTN mutant produced a large plaque morphology (**Extended Data Fig. 1a-b**), as seen with a FCS knockout mutant (ΔPRRA) ^2^. We then evaluated replication kinetics of ΔQTQTN in Vero E6 cells and found no difference between WT and ΔQTQTN (**Fig. 1d**). However, following direct 1:1 competition infections, the ΔQTQTN mutant had a significant advantage relative to WT SARS-CoV-2 in Vero E6 cells (**Fig. 1e**). This fitness advantage for ΔQTQTN likely explains the accumulation of this mutation in Vero E6-amplified virus stocks, as we also observed when this mutation emerges in Vero E6 cells infected with WT alone (**Extended Data Fig. 1c-d**). Notably, in Calu-3 2B4 cells, a human respiratory cell line, we observed a ~2.5 log reduction in ΔQTQTN replication at both 24 and 48 hours post infection (hpi) (**Fig. 1f**). Together, the results indicate that ΔQTQTN mutant is attenuated in respiratory cells and has a fitness advantage in Vero E6 cells; these results are similar findings to those we reported for the SARS-CoV-2 FCS knockout virus ^2^.

### ΔQTQTN attenuates disease, but not replication *in vivo*

We next evaluated the role of ΔQTQTN on virulence in an *in vivo* model. Three- to four-week-old male golden Syrian hamsters, which develop disease similar to that seen in humans ^12^, were intranasally inoculated with 10^5^ plaque-forming units (pfu) of WT SARS-CoV-2 or ΔQTQTN mutant and monitored for 7 days post infection (dpi) (**Fig. 2a**). Hamsters infected with WT steadily lost weight from 2 dpi with average peak weight loss of ~10% before beginning to recover at 5 dpi and regaining their starting weight by 7 dpi (**Fig. 2b**). The disease score peaks corresponded with maximum weight loss, with hamsters exhibiting ruffled fur, hunched posture, and/or reduced activity requiring additional welfare checks (**Fig. 2c**). Of note, 50% of WT-infected hamsters reached euthanasia criteria by 4 dpi (**Extended Data Fig. 2a**). In contrast, hamsters infected with ΔQTQTN experienced minimal weight loss and gained weight over the course of the infection (**Fig. 2b**). Similarly, no obvious disease was observed in ΔQTQTN infected animals (**Fig. 2c**). Hamsters infected with ΔQTQTN developed pulmonary lesions that were less extensive than those in hamsters infected with WT SARS CoV-2, involving smaller portions of the infected lungs on both days 2 and 4 after intranasal inoculation (**Fig. 2d**). All of the lesions were similar, with interstitial pneumonia, peribronchitis, peribronchiolitis, and vasculitis with predominantly subendothelial and perivascular infiltration by lymphocytes, and perivascular edema. Characteristic cytopathologic effects were observed in alveolar pneumocytes and bronchiolar epithelium, including cellular enlargement, binucleation and multinucleation, and prominent nucleoli. Together, the results demonstrate that the deletion of QTQTN motif attenuates SARS-CoV-2 disease *in vivo*.

**Figure 2:**
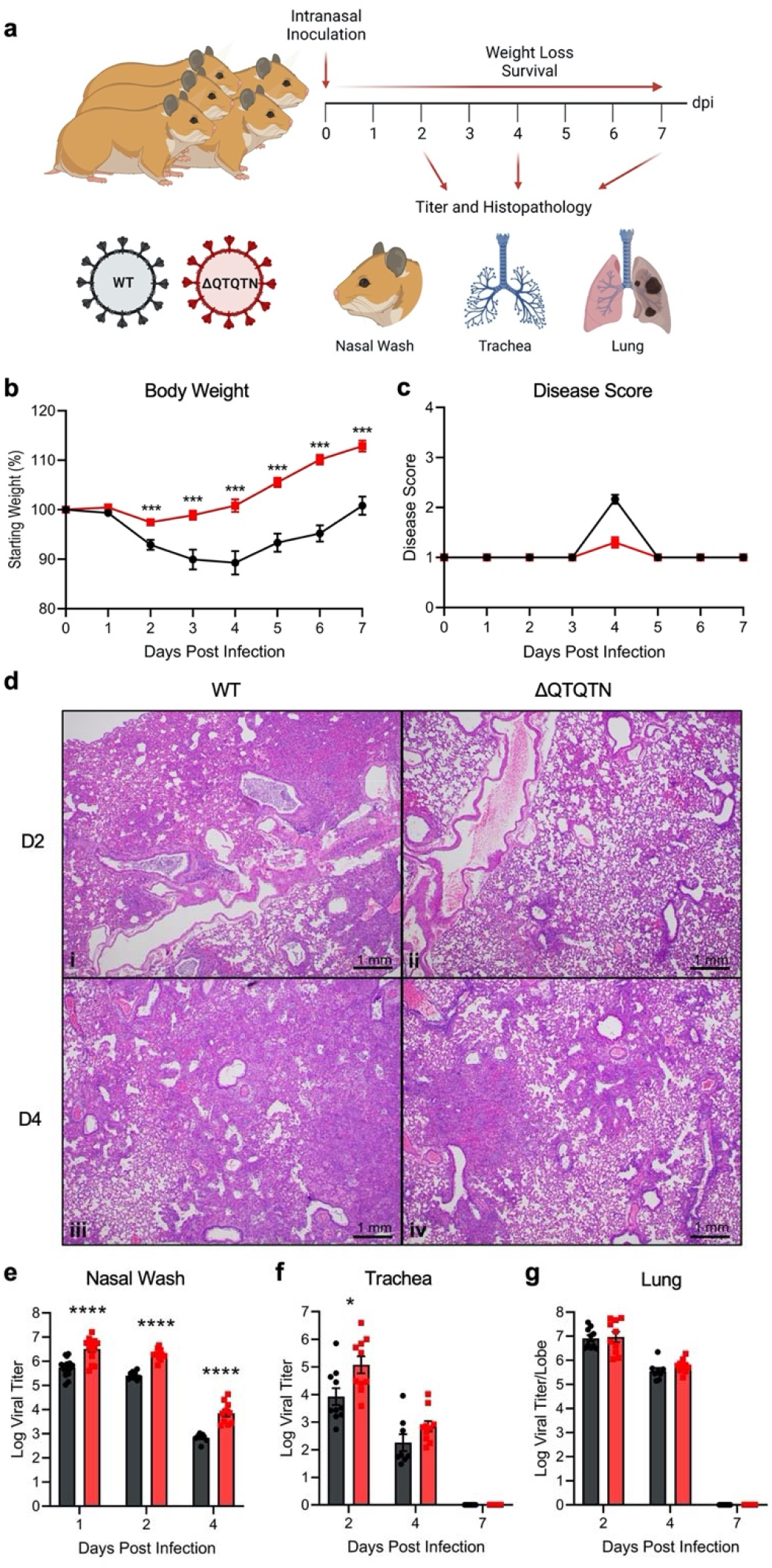
*In vivo* characterization of SARS-CoV-2 ΔQTQTN in golden Syrian hamsters. **a**, Schematic of golden Syrian hamster infection with WT (black) or ΔQTQTN (red) SARS-CoV-2. **b-c**, Three- to four-week-old male hamsters were infected with 10^5^ plaque-forming units (pfu) of WT or ΔQTQTN SARS-CoV-2 and monitored for weight loss (**b**) and disease score (**c**) for seven days (n=10). **d**, Histopathology of hamster lungs manifested more extensive lesions in animals infected with WT SARS-CoV-2 on day 2 (i) (4X magnification) than in animals infected with ΔQTQTN (ii) (4X). Lesions increased in volume on day 4 with greater proportions of the lungs affected in hamsters infected with WT (iii) (4X) than ΔQTQTN (iv) (4X) on day 4. **e-g**, Viral titers were measured for nasal washes (**e**), tracheae (**f**), and lungs (**g**). Data are mean ± s.e.m. Statistical analysis measured by two-tailed Student’s *t*-test. *, p≤0.05; **, p≤0.01; ***, p≤0.001; ****, p≤0.0001. Figures were created with BioRender.com

Despite clear attenuation in disease, ΔQTQTN viral replication in vivo was not compromised compared to WT SARS-CoV-2. In fact, viral titers were greater than WT SARS-CoV-2 with a 10-fold increase in nasal wash titers at 1, 2, and 4 dpi (**Fig. 2e**). Similar titer increases were observed in the trachea of infected hamsters at 2 dpi with equivalent titers at 4 dpi (**Fig. 2f**). Notably, viral titers were equivalent in the lungs for both 2 and 4 dpi (**Fig. 2g**). Examining viral RNA, we found that ΔQTQTN also had equivalent levels of viral replication relative to control SARS-CoV-2 in the lung (**Extended Data Fig. 2a**.). RNA expression data from hamster lung samples revealed clustering of WT and ΔQTQTN at 2 dpi and 4 dpi (**Extended Data Fig. 2b**). Of note, although more variability was present at 2 dpi, ΔQTQTN was slightly closer to mock samples at both time points. However, upregulated genes were similar between WT and ΔQTQTN in comparison to mock at both time points (**Extended Data Fig. 2c-d**). Together, these results indicate that attenuation of ΔQTQTN *in vivo* is not due to change in replication capacity. In addition, these data are consistent with *in vivo* results with the FCS knockout virus ^2^.

### ΔQTQTN reduces spike processing and entry

To examine the role of the QTQTN motif in spike processing, Vero E6 and Calu3-2B4 cells were infected with WT or ΔQTQTN and supernatant harvested at 24 hpi. Virus was then purified through sucrose cushion ultracentrifugation. Western blotting of the purified virus revealed reduced spike processing at the S1/S2 cleavage site for ΔQTQTN compared to WT in Vero E6 cells (**Fig. 3a**). Loss of the QTQTN motif resulted in little S1/S2 cleavage product and a significant increase in full-length spike compared to WT control. A similar reduction in spike processing was seen in Calu3-2B4 cells, although with more processing overall compared to in Vero E6 (**Fig. 3b**). Thus, deletion of the QTQTN motif impairs spike cleavage at the S1/S2 site, similar to findings with the SARS-CoV-2 mutants lacking the FCS ^2,7^.

**Figure 3:**
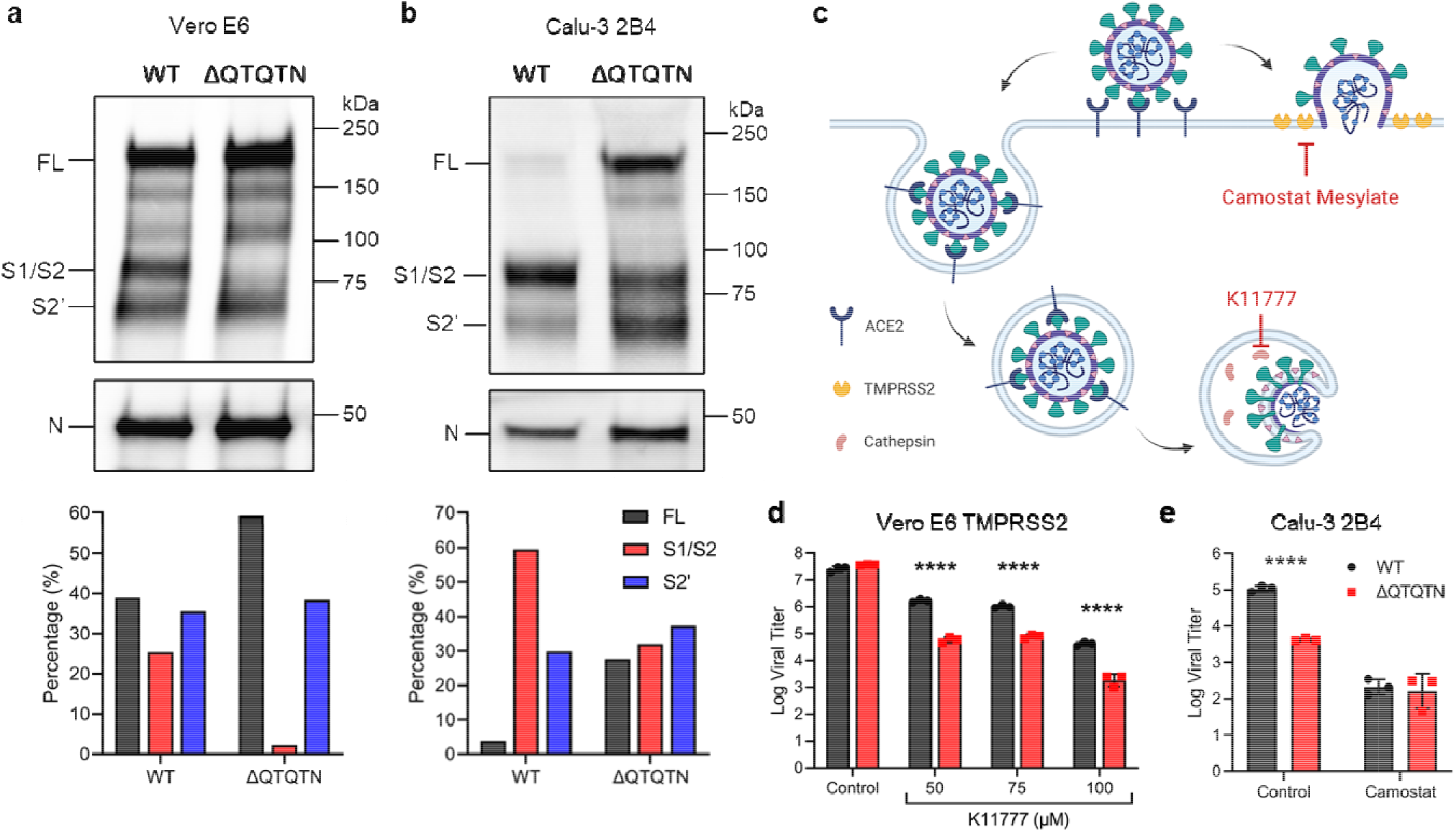
QTQTN motif is involved in spike processing and protease usage. **a-b**, Purified WT and ΔQTQTN SARS-CoV-2 virions from Vero E6 (**a**) and Calu-3 2B4 (**b**) cells probed with anti-S or anti-N antibody (upper). Full-length (FL), S1/S2 cleavage product, and S2’ cleavage product are indicated. Quantification of densitometry of FL (black), S1/S2 (red), and S2’ (blue) normalized to N SARS-CoV-2 protein shown (lower). Results are representative of two experiments. **c**, Schematic of SARS-CoV-2 entry and use of host proteases. Inhibitors for TMRPSS2 (camostat mesylate) and cathepsin (K11777) are indicated. **d**, Viral titers at 24 hpi from TMPRSS2-expressing Vero E6 cells pretreated with varying doses of cathepsin inhibitor K11777 and infected with WT (black) or ΔQTQTN SARS-CoV-2 (red) at an MOI of 0.01 pfu/cell (n=3). **e**, Viral titer at 24hpi from Calu-3 2B4 pretreated with 100 μM of camostat mesylate and infected with WT or ΔQTQTN SARS-CoV-2 at an MOI of 0.01 (n=3). Data are mean ± s.d. Statistical analysis measured by two-tailed Student’s *t*-test. *, p≤0.05; **, p≤0.01; ***, p≤0.001; ****, p≤0.0001. Figures were created with BioRender.com

After receptor binding, the CoV spike protein is cleaved by a host protease as part of the virus entry process. Different proteases can be utilized to activate the spike fusion machinery: serine proteases like TMPRSS2 at the cell surface or cathepsins within endosome (**Fig. 3c**). Our prior work found that the absence of TMPRSS2 in Vero E6 cells plays a role in selection of SARS-CoV-2 strains with FCS deletions ^2^. Calu-3 2B4 cells also express high levels of TMPRSS2. We therefore hypothesized that the absence of TMPRSS2 activity contributes to ΔQTQTN selection in Vero E6 and attenuation in Calu3 2B-4 cells. To test this hypothesis, Vero E6 cells expressing TMPRSS2 were pretreated with cathepsin inhibitor K11777 before infection with WT or ΔQTQTN, and viral titers were measured at 24 hpi. With cathepsin inhibited and TMPRSS2 activity intact, a significant, ~1.5 log reduction in viral titer was observed for ΔQTQTN compared to WT over a dose range of K11777, mirroring the attenuation observed in the Calu-3 2B4 cells (**Fig. 3d**). Infection of untreated TMPRSS2-expressing Vero E6 revealed no difference in replication between WT and ΔQTQTN (**Extended Data Fig. 3**). When Calu-3 2B4 cells were pretreated with the serine protease inhibitor camostat mesylate, WT SARS-CoV-2 titers were reduced and equivalent to ΔQTQTN (**Fig. 3e**). Together, these data indicate that the loss of the QTQTN motif reduces the capacity of the virus to use TMPRSS2 for entry.

### Glycosylation of the QTQTN motif contributes to spike processing

As the absence of the QTQTN motif attenuates SARS-CoV-2, we set out to determine if the QTQTN motif itself has a significant role during infection. Notably, the second threonine, T678, of the motif has been previously shown to be O-linked glycosylated ^4^. Structurally, the QTQTN site resides on an exterior loop of the spike and is capable of accommodating large glycans, which may contribute to interactions with proteases like TMPRSS2 (**Fig. 4a**). To determine the role of glycosylation in spike processing, we generated mutants abolishing the glycosylated T678 (QTQVN) alone or together with the first threonine T676 (QVQVN) to exclude possible compensatory glycosylation (**Fig. 4b**). Similar to ΔQTQTN, the glycosylation mutations did not affect virus yield with titers comparable to WT (**Extended Data Fig. 4a**). However, plaque morphologies of QTQVN and QVQVN were more similar to WT than to the ΔQTQTN mutant (**Extended Data Fig. 4b**). Viral replication in Vero E6 cells was not affected for QTQVN and QVQVN mutants; however, both were attenuated at 24 hpi in Calu-3 2B4 cells, mirroring what was observed with the ΔQTQTN mutant (**Fig. 4c-d**). Moreover, viral titers of the glycosylation mutants reached WT levels by 48 hpi in Calu-3 2B4. Together, these results indicate that the loss of glycosylation sites in the QTQTN motif attenuates replication in Calu-3 2B4 cells.

**Figure 4:**
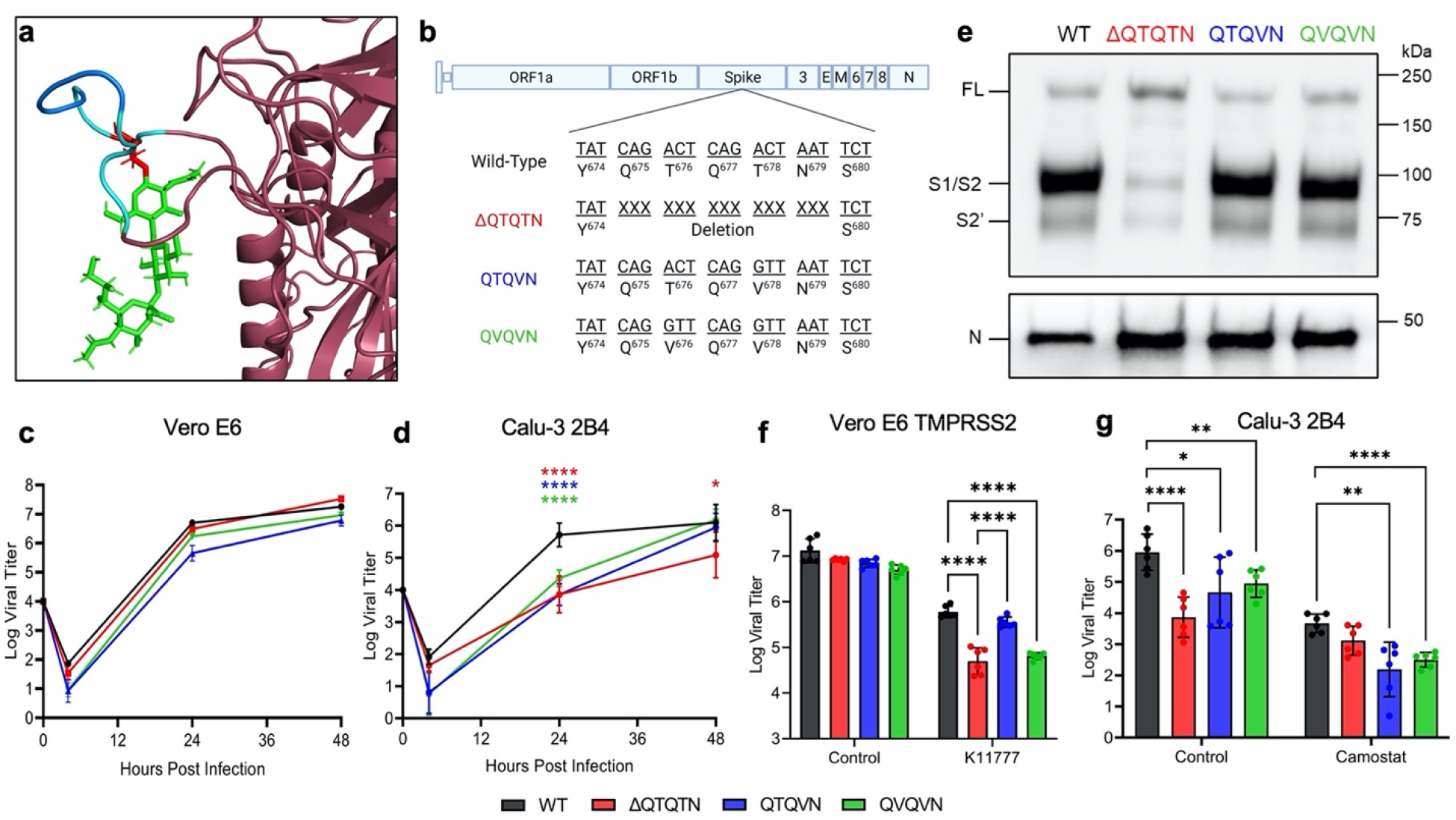
Glycosylation of QTQTN motif contributes to SARS-CoV-2 virulence. **a**, Structural modeling of O-linked glycosylation on threonine 678 (red) of QTQTN motif. PRRA (blue) remains exposed. **b**, Schematic of SARS-CoV-2 genome with glycosylation mutations. **c**, Viral titers from Vero E6 cells infected with WT (black), ΔQTQTN (red), QTQVN (blue), or QVQVN (green) SARS-CoV-2 at an MOI of 0.01 (n=3). **d**, Viral titers from Calu-3 2B4 cells infected with WT, ΔQTQTN, QTQVN, or QVQVN SARS-CoV-2 at an MOI of 0.01 (n=6). **e**, Purified WT, ΔQTQTN, QTQVN, and QVQVN SARS-CoV-2 virions from Calu-3 2B4 probed with anti-S or anti-N antibody. Full-length (FL), S1/S2 cleavage product, and S2’ cleavage products are indicated. Results are representative of two experiments. **f**, Viral titers at 24 hpi from TMPRSS2-expressing Vero E6 cells pretreated with 50 μM of K11777 and infected with WT, ΔQTQTN, QTQVN, or QVQVN SARS-CoV-2 at an MOI of 0.01 (n=6). **g**, Viral titers at 24hpi from Calu-3 2B4 pretreated with 50 μM of camostat mesylate and infected with WT, ΔQTQTN, QTQVN, or QVQVN SARS-CoV-2 at an MOI of 0.01 (n=6). Data are mean ± s.d. Statistical analysis measured by two-tailed Student’s *t*-test relative to WT. *, p≤0.05; **, p≤0.01; ***, p≤0.001; ****, p≤0.0001.

We next examined if the glycosylation-defective mutants had altered spike processing similar to ΔQTQTN. Western blotting of virus purified from Calu-3 2B4 cells by sucrose cushion ultracentrifugation revealed intact spike processing at the S1/S2 site with both glycosylation mutants (**Fig. 4e, Extended Data Fig. 4c**). Contrasting with ΔQTQTN, the QTQVN and QVQVN mutants had significant S1/S2 spike cleavage with levels similar to WT SARS-CoV-2. Likewise, spike processing of QTQVN and QVQVN in normal and TMPRSS2-expressing Vero E6 cells was similar to that of WT (**Extended Data Fig. 4d-e)**. Together, these results suggest that the loss of glycosylated residues does not impact spike processing of SARS-CoV-2.

To further understand the role of glycosylation in spike processing, we next examined if glycosylation is involved in protease interaction and usage. Using TMPRSS2-expressing Vero E6, we pretreated cells with K11777 to disrupt cathepsin activity, infected with WT, ΔQTQTN, QTQVN, or QVQVN, and examined viral titers at 24 hpi (**Fig. 4f**). While the ΔQTQTN titer was reduced compared to WT as observed before, the QTQVN titer was equivalent to WT (**Fig. 4f**). In contrast, disruption of both glycosylation residues with the QVQVN mutant titer resulted in attenuation equivalent to that of ΔQTQTN, suggesting that abolishing both O-linked glycosylation sites disrupted TMPRSS2 utilization (**Fig. 4f**). We subsequently pretreated Calu-3 2B4 cells with camostat mesylate to disrupt serine protease activity and infected with the glycosylation mutants (**Fig. 4g**). Interestingly, while treatment with camostat in Calu-3 2B4 cells reduced WT titer to equivalent levels as ΔQTQTN, viral titers of QTQVN and QVQVN mutants were even lower. However, the overall differences in titer between WT and the glycosylation mutants was reduced, suggesting that glycosylation is important for TMPRSS2 utilization and entry (**Fig. 4g**). Overall, these results argue that glycosylation of the QTQTN motif is important to protease interactions with spike and SARS-CoV-2 infection.

### The QTQTN motif is necessary for efficient SARS-CoV-2 infection and pathogenesis

The presence of the FCS in SARS-CoV-2 plays a critical role in infection and pathogenesis by facilitating an increase spike processing upon nascent virion release ^2^. This FCS, unusual to SARS-like coronaviruses, has been highlighted as a potential “smoking gun” for an engineered virus ^13^. Yet, the FCS alone is insufficient to drive infection and pathogenesis. The upstream QTQTN motif adds two distinct elements that contribute to this capacity and virulence. The loss of the QTQTN motif produces a shorter, more rigid exterior loop in the spike, likely reducing access to the FCS. The result is a significant reduction in spike processing and attenuation of the ΔQTQTN mutant both *in vitro* and *in vivo*. Similarly, while mutations of the glycosylation residues in the QTQTN motif do not change overall spike processing, the modification of the motif attenuates virus replication in a TMPRSS2-dependent manner. Overall, our results argue that the FCS, the length/composition of the exterior loop, and glycosylation of the QTQTN motif are all needed for efficient infection and pathogenesis (**Figure 5**). Disruption of any of these three elements attenuates SARS-CoV-2, highlighting the complexity of spike activation beyond the simple presence of a furin cleavage site.

**Figure 5:**
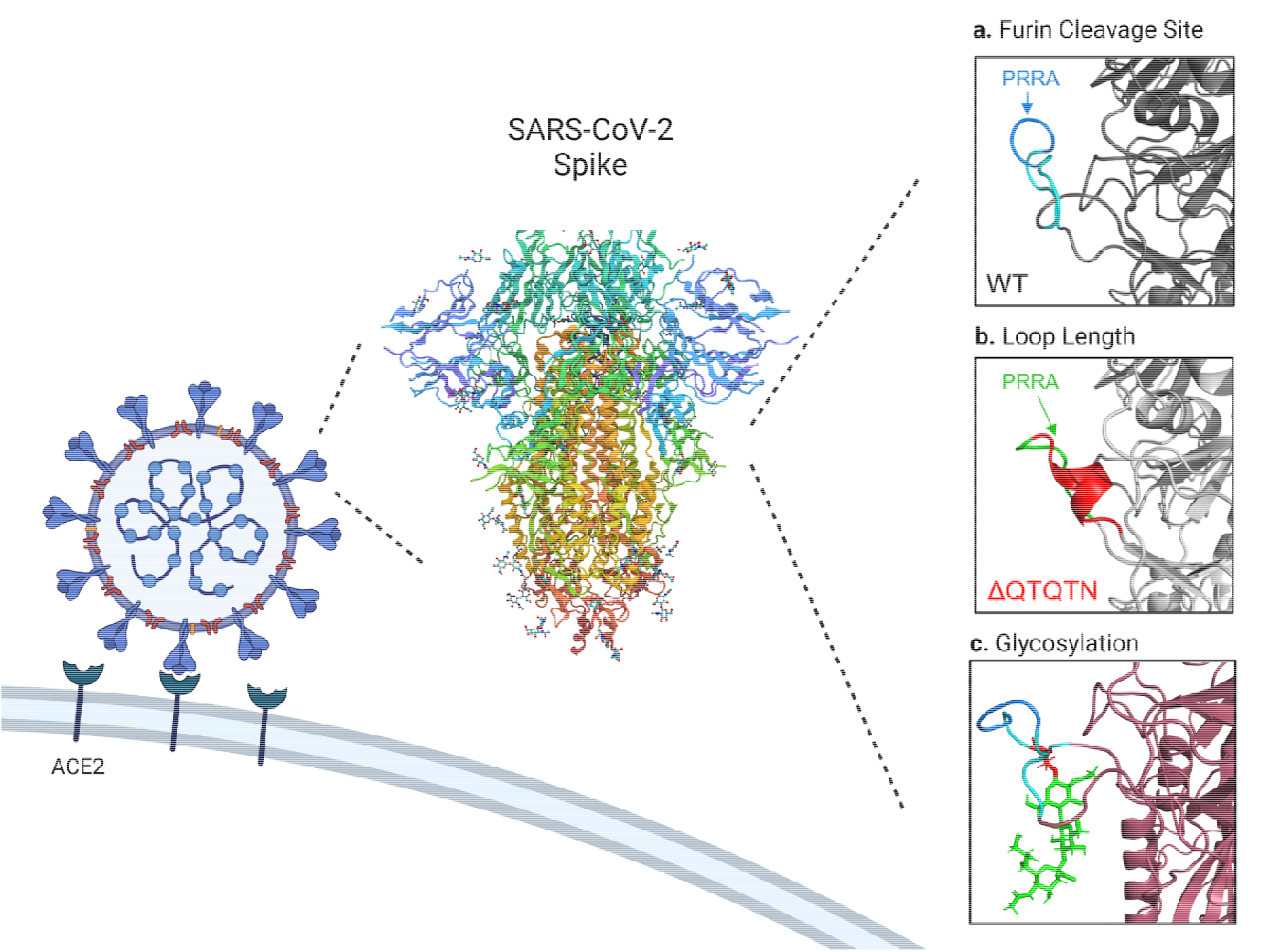
Characteristics of SARS-CoV-2 S1/S2 spike cleavage site for efficient infection. The SARS-CoV-2 S1 spike cleavage site contains multiple components required for efficient infection and virulence: the furin cleavage site (FCS), PRRA, is important for spike processing (**a**); the loop length/composition, affected by the QTQTN motif, manages FCS accessibility and protease interaction (**b**); and glycosylation is also involved in the protease interaction (**c**). SARS-CoV-2 spike^14^. Figures were created with BioRender.com

## Acknowledgments

We would like to thank Shinji Makino for gifting the nucleocapsid antibody. Figures were created with BioRender.com

## Author contributions

Conceptualization: VDM

Formal analysis: MNV, JAP, ALR, VDM

Funding acquisition: SCW, ALR, VDM

Investigation: MNV, KGL, JAP, DS, BAJ, SS, DS, CS, REA, PACV, DHW, KD

Methodology: MNV, ALR, KSP, VDM, SCW

Project Administration: VDM

Supervision: SCW, DHW, ALR, KSP, VDM

Visualization: MNV, KD, DHW, VDM

Writing – original draft: MNV, VDM

Writing – review and editing: MNV, VDM, SCW

---

**Extended Data Fig. 1:**
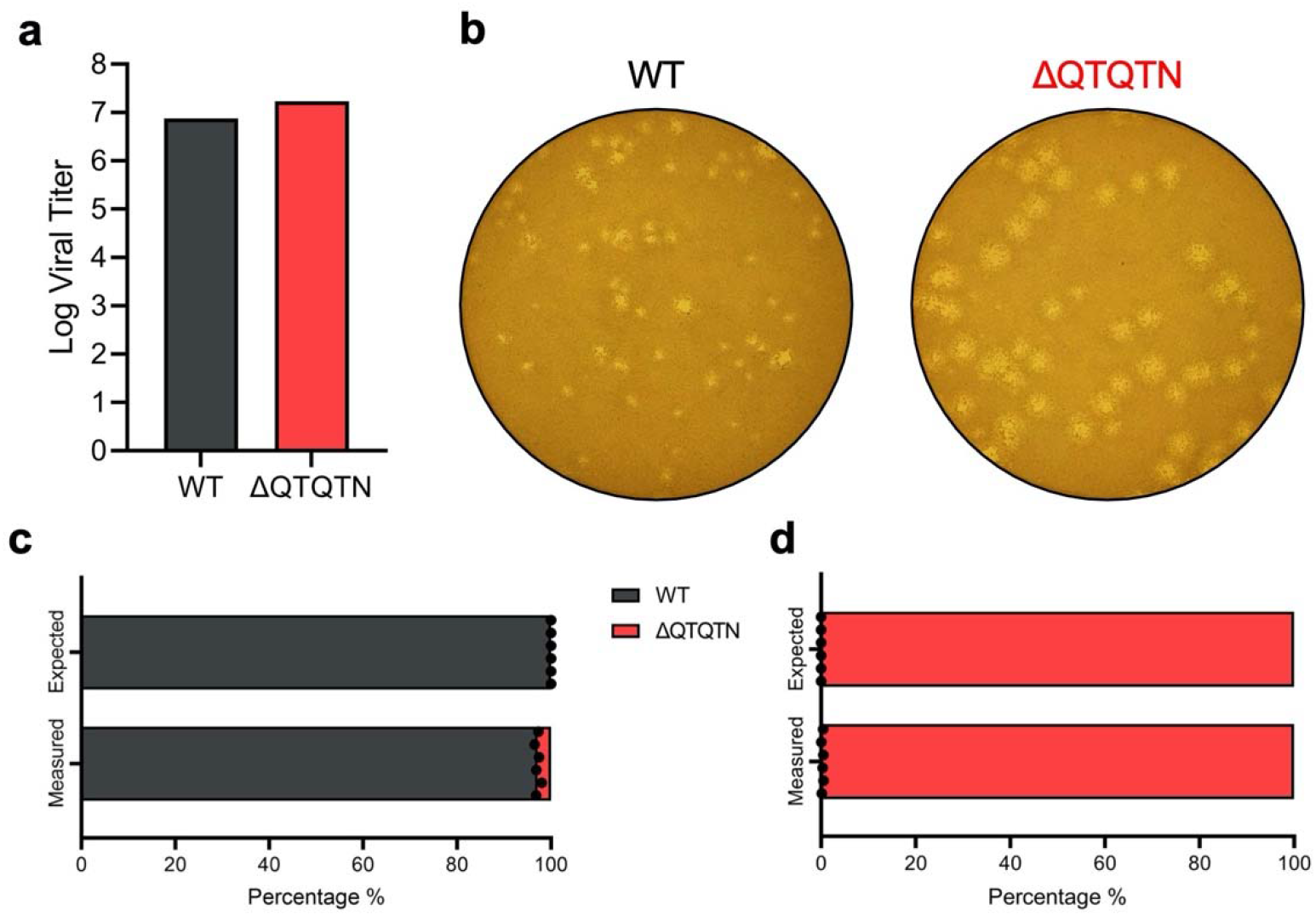
ΔQTQTN SARS-CoV-2 replication. **a**, Virus stock titer of WT and ΔQTQTN SARS-CoV-2 from Vero E6. **b**, Plaque morphology of WT and ΔQTQTN in Vero E6. **c-d**, Competition assay between WT and ΔQTQTN SARS-CoV-2 at a ratio of 100:0 (**c**) and 0:100 (**d**) WT:ΔQTQTN, showing RNA percentage from next generation sequencing.

**Extended Data Fig. 2:**
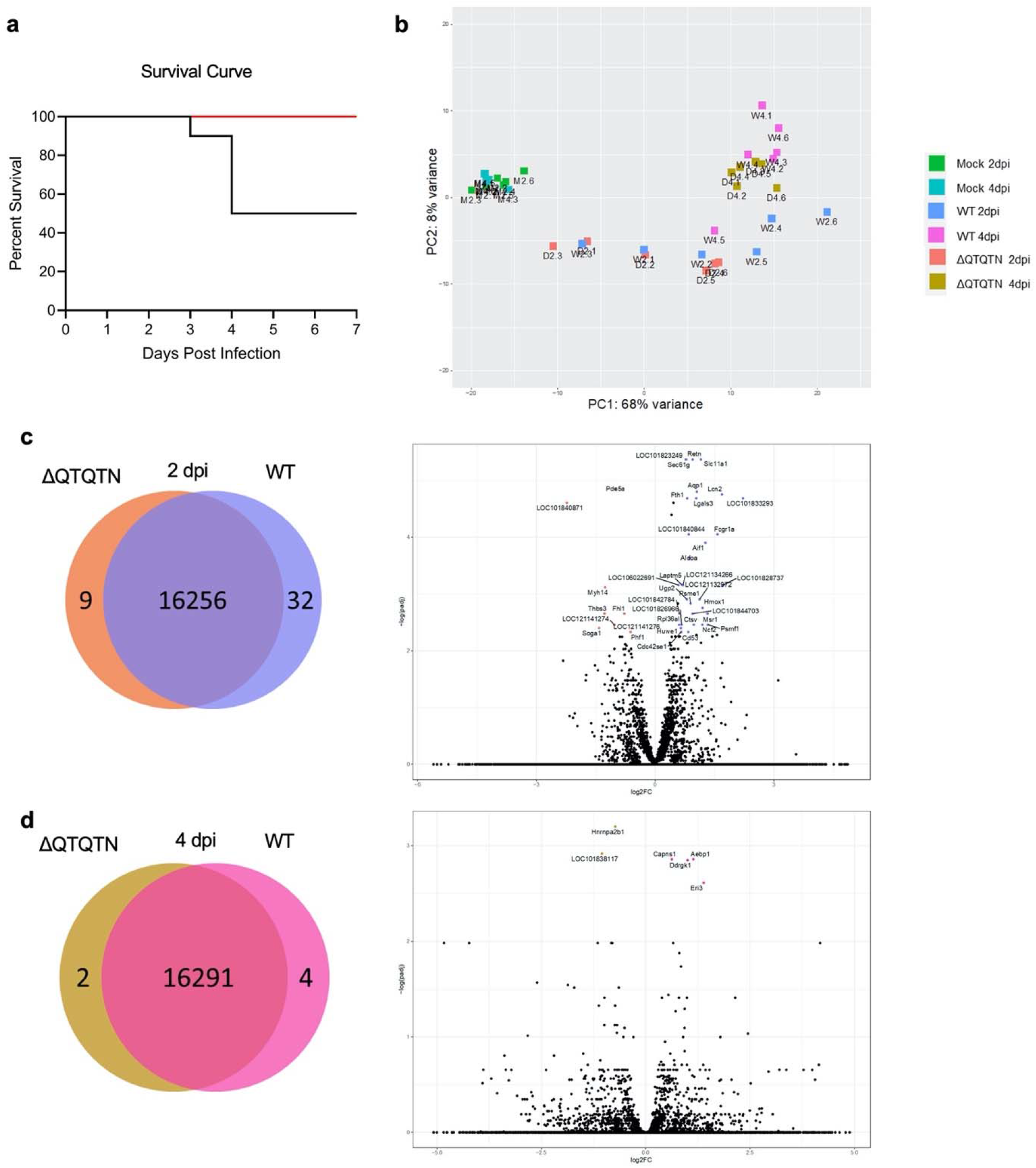
Hamster infection with ΔQTQTN SARS-CoV-2. **a**, Survival curve (based on euthanasia criteria of >20% weight loss) following infection of WT or ΔQTQTN SARS-CoV-2 (n=10). **b**, Principal component analysis (PCA) plot of hamster lung samples. **c**, DESeq2 analysis of mapped genes between WT (purple) and ΔQTQTN (orange) at 2 dpi (left) with upregulated genes indicated in volcano plot (right). **d**, DESeq2 analysis of mapped genes between WT (purple) and ΔQTQTN (orange) at 4 dpi (left) with upregulated genes indicated in volcano plot (right).

**Extended Data Fig. 3:**
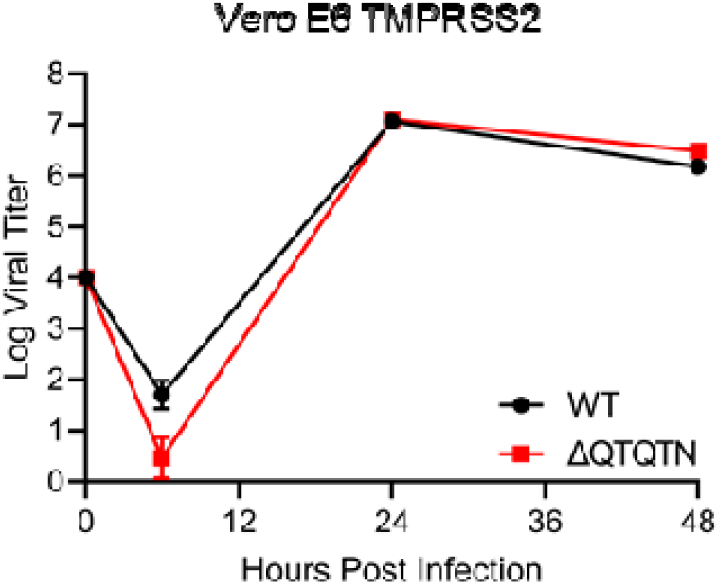
ΔQTQTN SARS-CoV-2 replication in TMPRSS2-expressing Vero E6. Viral titer from TMPRSS2-expressing Vero E6 infected with WT or ΔQTQTN SARS-CoV-2 at an MOI of 0.01 (n=3). Data are mean ± s.d. Statistical analysis measured by two-tailed Student’s *t*-test. *, p≤0.05; **, p≤0.01; ***, p≤0.001; ****, p≤0.0001.

**Extended Data Fig. 4:**
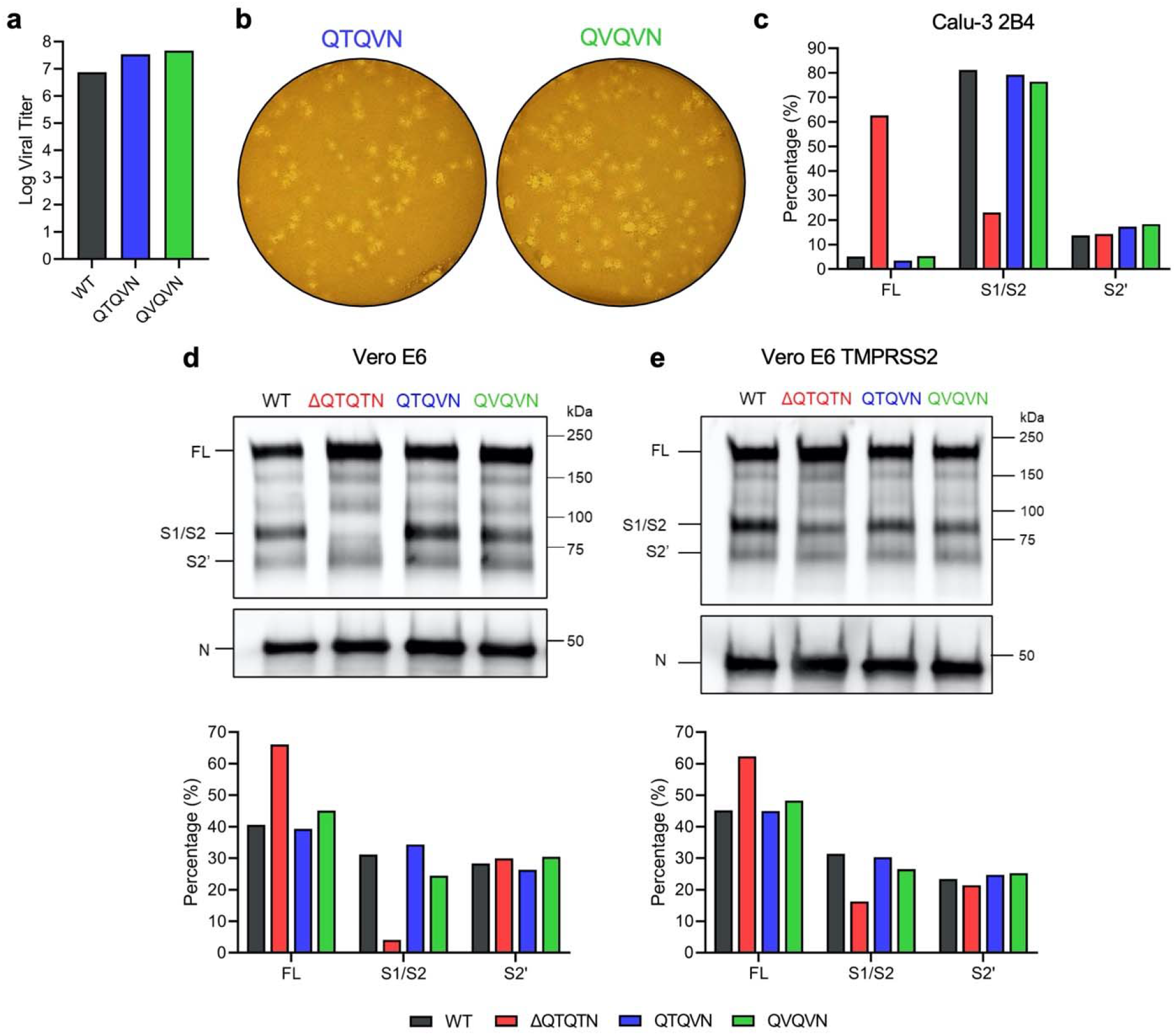
QTQVN and QVQVN SARS-CoV-2 replication and spike processing. **a**, Virus stock titer of WT, QTQVN, and QVQVN SARS-CoV-2 in Vero E6. **b**, Plaque morphology of QTQVN and QVQVN in Vero E6. **c**, Quantification by densitometry of full-length (FL), S1/S2 cleavage product, and S2’ cleavage product from western blot experiments of glycosylation mutants in Calu-3 2B4. **d-e**, Purified WT (black), ΔQTQTN (red), QTQVN (blue), and QVQVN (green) SARS-CoV-2 virions from Vero E6 (**d**) and TMRPSS2-expressing Vero E6 (**e**) probed with anti-S or anti-N antibody (upper). Full-length (FL), S1/S2 cleavage product, and S2’ cleavage product are indicated. Quantification of densitometry of FL, S1/S2, and S2’ normalized to N shown (lower). Results are representative of two experiments.

## Methods

### Cell culture

Vero E6 cells were grown in DMEM (Gibco #11965-092) supplemented with 10% fetal bovine serum (FBS) (HyClone #SH30071.03) and 1% antibiotic-antimycotic (Gibco #5240062). Calu-3 2B4 cells were grown in DMEM supplemented with 10% defined FBS (HyClone #SH30070.03), 1% antibiotic-antimycotic, and 1 mg/ml sodium pyruvate. Vero E6 expressing TMPRSS2 cells were grown in DMEM (Gibco #11885-084) supplemented with 10% FBS and 1 mg/ml geneticin (Gibco #10131035).

### Viruses

The recombinant wild-type (WT) and mutant SARS-CoV-2 virus sequences are based on the USA-WA1/2020 isolate sequence provided by the World Reference Center for Emerging Viruses and Arboviruses (WRCEVA), which was originally obtained from the USA Centers for Disease Control and Prevention (CDC) as previously described ^15^. The mutant viruses (ΔQTQTN, QTQVN, and QVQVN) were generated using standard cloning techniques and reverse genetics system as previously described ^2,10^. Standard plaque assays were used for virus titer.

### *In vitro* infection

Viral infections in Vero E6, Calu-3 2B4 and TMRPSS2-expressing Vero E6 cells were performed as previously described ^11,16^. Briefly, cells were washed with PBS and infected with WT or mutant SARS-CoV-2 at an MOI of 0.01 for 45 min at 37°C. Following absorption, cells were washed three times with PBS and fresh growth media was added. Three or more biological replicates were collected at each time point.

### Protease inhibitor treatment

TMPRSS2-expressing Vero E6 or Calu-3 2B4 cells were pretreated with 50-100 μM of K11777 (AdipoGen #AG-CR1-0158-M005) or 50-100 μM of camostat mesylate (Sigma-Aldrich #SML0057-10MG), respectively, in 1 ml growth medium for 1 hr at 37°C. Cells were subsequently washed with PBS and infected with WT or mutant SARS-CoV-2 at an MOI of 0.01 as described in ‘*In vitro* infection’.

### Competition assay and next generation sequencing analysis

Ratios (1:0, 1:1, and 0:1 WT:ΔQTQTN) for the competition assays were determined by pfu of virus stock. Vero E6 cells were infected with a total MOI of 0.01 (WT alone, 1:1 WT:ΔQTQTN, or ΔQTQTN alone) as described in ‘*In vitro* infection’. RNA was collected from cell lysate with Trizol reagent (Invitrogen #15596018) and extracted with Direct-zol RNA Miniprep Plus kit (Zymo #R2072). RNA libraries were prepared by ClickSeq and sequenced as previously described ^2,17^.

### Virion purification and western blotting

Vero E6, Calu-3 2B4 and TMPRSS2-expressing Vero E6 cells were infected with WT or mutant SARS-CoV-2 at an MOI of 0.01. Culture supernatant was harvested 24 hpi and clarified by low-speed centrifugation. Virus particles were then pelleted by ultracentrifugation through a 20% sucrose cushion at 26,000 rpm for 3 hr using a Beckman SW28 rotor. Pellets were resuspended in 2x Laemmli buffer to obtain protein lysates. Relative viral protein levels were determined by SDS–PAGE followed by western blot analysis as previously described ^2,15,18,19^. In brief, sucrose-purified WT and mutant SARS-CoV-2 virions were inactivated by boiling in Laemmeli buffer. Samples were loaded in equal volumes into 4–20% Mini-PROTEAN TGX Gels (Bio-Rad #4561093) and electrophoresed by SDS–PAGE. Protein was transferred to polyvinylidene difluoride (PVDF) membranes. Membranes were probed with SARS-CoV S-specific antibodies (Novus Biologicals #NB100-56578) and followed with horseradish peroxidase (HRP)-conjugated anti-rabbit antibody (Cell Signaling Technology #7074S). Membranes were stripped and reprobed with SARS-CoV N-specific antibodies (provided by S. Makino) and the HRP-conjugated anti-rabbit secondary IgG to measure loading. Signal was developed using Clarity Western ECL substrate (Bio-Rad #1705060) or Clarity Max Western ECL substrate (Bio-Rad #1705062) and imaging on a ChemiDoc MP System (Bio-Rad #12003154). Densitometry was performed using ImageLab 6.0.1 (Bio-Rad #2012931).

### Hamster infection studies

Male golden Syrian hamsters (3-4 weeks old) were purchased from Envigo. All studies were conducted under a protocol approved by the UTMB Institutional Animal Care and Use Committee and complied with USDA guidelines in a laboratory accredited by the Association for Assessment and Accreditation of Laboratory Animal Care. Procedures involving infectious SARS-CoV-2 were performed in the Galveston National Laboratory ABSL3 facility. Animals were housed in groups of five and intranasally inoculated with 10^5^ pfu of WT or ΔQTQTN SARS-CoV-2. Animals were monitored daily for weight loss and development of clinical disease through the course of the study. Hamsters were anesthetized with isoflurane (Henry Schein Animal Health) for viral infection and nasal washes.

### Histology

Left lungs were harvested from hamsters and fixed in 10% buffered formalin solution for at least 7 days. Fixed tissue was then embedded in paraffin, cut into 5 μM sections, and stained with hematoxylin and eosin (H&E) on a SAKURA VIP6 processor by the University of Texas Medical Branch Surgical Pathology Laboratory.

### Structural modeling

Structural models were generated using SWISS-Model to generate homology models for WT, ΔQTQTN, and glycosylated QTQTN SARS-CoV-2 spike protein on the basis of the SARS-CoV-1 trimer structure (Protein Data Bank code 6ACD). Homology models were visualized and manipulated in MacPyMol (version 1.3).

### Transcriptomics

Hamster lungs were homogenized in Trizol reagent (Thermo Fisher) and RNA was extracted with Direct-zol RNA Miniprep Plus kit (Zymo #R2072). The short-read sequencing libraries were generated from extracted RNA using Poly-A Click-Seq (PAC-Seq) ^20,21^. Briefly, RNAs containing poly(A) tails were selectively reverse transcribed with oligo(dT) primers and stochastically terminated with azido-NTPS. Libraries were then gel purified (200-400 bp) and sequenced using Illumina platform (NextSeq550). Differential Poly-A Cluster (DPAC) was used to identify changes in overall expression ^20,21^. An p-adjusted value (p-adj) of <0.1 and an absolute value of log2 fold change (|log2FC|) greater than 0.58 (or minimum of 50% increase/decrease) was used to filter results. The command ran for this data set was ~/DPAC -p PMCDB -t 4 -x [flattened_annotations] -y [reference_names] -g [genome] -n 6 -v golden_hamster,Mesaur [metadata_file] [index] [experiment name] [output_directory], where -p indicates parameters used, in this case P (perform data pre-processing), M (map data), C (force new PAS cluster generation), D (perform differential APA analysis), and B (make individual bedgraphs), -t indicates how many threads were to be used, and -n indicates number of replicates. Annotations, gene names, index were used for Syrian golden hamster and mapped to the Syrian golden hamster genome (Mesaur). Data quality was determined to be sufficient by generating and loading bedgraph files into the UCSC Genome Browser ^22^.

